# Identification of evolutionarily conserved nuclear matrix proteins and their prokaryotic origins

**DOI:** 10.1101/863969

**Authors:** Rahul Sureka, Rakesh Mishra

## Abstract

Compared to prokaryotic cells, a typical eukaryotic cell is much more complex along with its endomembrane system and membrane-bound organelles. Although the endosymbiosis theories convincingly explain the evolution of membrane-bound organelles such as mitochondria and chloroplasts, very little is understood about the evolutionary origins of the nucleus, the defining feature of eukaryotes. Most studies on nuclear evolution have not been able to take into consideration the underlying structural framework of the nucleus, attributed to the nuclear matrix (NuMat), a ribonucleoproteinaceous structure. This can largely be attributed to the lack of annotation of its core components. Since, NuMat has been shown to provide a structural platform for facilitating a variety of nuclear functions such as replication, transcription, and splicing, it is important to identify its protein components to better understand these processes. In this study, we address this issue using the developing embryos of *D. melanogaster* and *D. rerio* and identify 362 core NuMat proteins that are conserved between the two organisms. We find that of them, 132 protein groups have originated from pre-existing proteins in prokaryotes. While 51 were conserved across all eukaryotic supergroups, 17 new proteins evolved before the evolution of the last eukaryotic common ancestor and together these 68 proteins out of the 362 core conserved NuMat proteins are conserved across all eukaryotes indicating their indispensable nature for nuclear function for over 1.5 billion years of eukaryotic history. Our analysis paves the way to understand the evolution of the complex internal nuclear architecture and its functions.

## Introduction

The origin of the eukaryotic cell ~1.5 billion years ago marked a step change in the evolutionary history of life on earth. It led to the diversification of a wide variety of lineages displaying different morphologies, several of which independently evolved into complex multicellular organisms (Dacks et al. 2016). Compared to an average prokaryotic cell, the eukaryotic cell exhibits a considerable increase in structural complexity, typified by the presence of an endomembrane system separating the nucleus from the rest of the cytoplasm and other membrane-bound organelles. There is compelling genomic evidence that the mitochondria and chloroplast arose from two primary endosymbiotic events: one from the endosymbiosis of an α-proteobacterium giving rise to the mitochondria, and the other from the endosymbiosis of a cyanobacterium giving rise to chloroplasts (Margulis 1970; Pace 2009). While the endosymbiotic theories for the evolution of other membrane-bound organelles such as mitochondria and chloroplast are generally accepted, very little is known and even less is agreed upon, about the events that led to the evolution of the nucleus, the quintessential feature of eukaryotes.

Our knowledge about the early evolution of nucleus, and by extension eukaryotes, remain unsatisfactory because of two reasons: Firstly, most studies on nuclear evolution focus on the nuclear membrane and the proteins associated with it. This is because the nuclear membrane is the most prominent and an undisputed part of nuclear morphology and its biochemical composition is relatively well documented, at least for model organisms (animals, plants and fungi) (Wilson and Dawson 2011). In contrast, the dynamic and complex internal architecture of the nucleus, first proposed to be supported by the nuclear matrix (NuMat) by Berezney and Coffey in 1974 (Berezney and Coffey 1974), has not been incorporated in studies on nuclear evolution. Although various studies over the years have shown that the NuMat provides a structural framework for the functional regulation of most, if not all, nuclear metabolic processes such as DNA replication, transcription, splicing, DNA repair, and higher order chromatin organization (Mirkovitch et al. 1984; Jackson and Cook 1985; Jackson, D. A. 1986; Zeitlin et al. 1987; Koehler and Hanawalt 1996), the in-vivo existence of this structure has been widely debated due to concerns about the artefactual attachment of nuclear components during preparation (Pederson 2000; Razin et al. 2014). The issue is further complicated by the fact that the biochemical composition of NuMat varies widely depending on extraction methods employed and has left the question of identifying the core components of NuMat unanswered (Nickerson 2001; Simon and Wilson 2011). This lack of annotation of the core nucleoskeletal proteins has led to the exclusion of NuMat composition from consideration in studies of nuclear evolution, impeding the progress of the field (Wilson and Dawson 2011). Secondly, in recent years, attempts have been made to circumvent this drawback by genome-wide comparisons of extant eukaryotes. However, this approach is limited because the genetic diversity of extant eukaryotes, particularly unicellular eukaryotes, is not yet reflected in the completed genome projects (D. Campo et al. 2014). Eukaryotes are divided into six major supergroups: Opisthokonts (e.g., fungi, animals, protists), Amoebozoa (e.g., Dictyostelium), Excavates (e.g., Trypanosomes, Giardia), Chromalveolates (e.g., Plasmodium), Archaeplastids (e.g., plants), and Rhizaria (e.g., Plasmodiophora) (Hampl et al. 2009), out of which two supergroups, i.e., Opisthokonts and Archaeplastids are overwhelmingly represented among the reasonably completed genome assemblies. Moreover, although genome-wide comparisons are useful for predicting genes constituting the last eukaryotic common ancestor (LECA), they have limited utility in predicting the subset of genes which were essential for the structural evolution of nucleus. Therefore, it is imperative to identify the core structural components of the nucleus which includes both the NuMat and the nuclear membrane.

In the present study, we identify the core NuMat proteins by comparing the NuMat proteomes from two different developmental stages of *Drosophila melanogaster* and *Danio rerio*. We then use this information to identify the prokaryotic lineages which contributed to the origins of these NuMat proteins, and also to identify the NuMat proteins which are conserved in organisms spanning across all eukaryotic supergroups. These findings present a way forward to understand the structural evolution of the nucleus and its associated functions.

## Results

### Experimental approach

In order to identify the core protein composition of NuMat, we used the developing embryos of *D. melanogaster* and *D. rerio* as our model system. The developing embryo gives a unique opportunity because we can study both undifferentiated cells in the early stages (pre-gastrulation stage) of development and several types of differentiated cells in the late embryonic stages from an *in vivo* source. We reasoned that since the NuMat components are known to be dynamic (Varma and Mishra 2011), comparing the NuMat proteome of undifferentiated and differentiated cells would help in identifying the constitutive as well as cell-type dependent protein constituents associated with it. Therefore, we used 0-2 hr (early stage with undifferentiated cells) and 14-16 hr embryos (late stage with differentiated cells) of *D. melanogaster* and four hours post fertilization (4 hpf) (early stage with undifferentiated cells) and one day post fertilization (1 dpf) embryos (late stage with differentiated cells) of *D. rerio* for our experiments.

### NuMat preparation and quality control

Pure nuclei were isolated from each of early and late embryos of *D. melanogaster* and *D. rerio* using sucrose gradient centrifugation and were found to be free of unlysed cells and other membranous contamination when observed under fluorescence microscope (Figure 1 A, C). The nuclei were also checked for cytosolic and mitochondrial contamination by western hybridization (Figure 1 B, D). The blots showed that while Lamin was present in both whole embryo lysates and isolated nuclei from both organisms, GAPDH, which is a cytosolic marker, and Cytochrome C, which is a mitochondrial marker, were present only in whole embryo lysates but not in pure nuclei.

**Figure 1:**
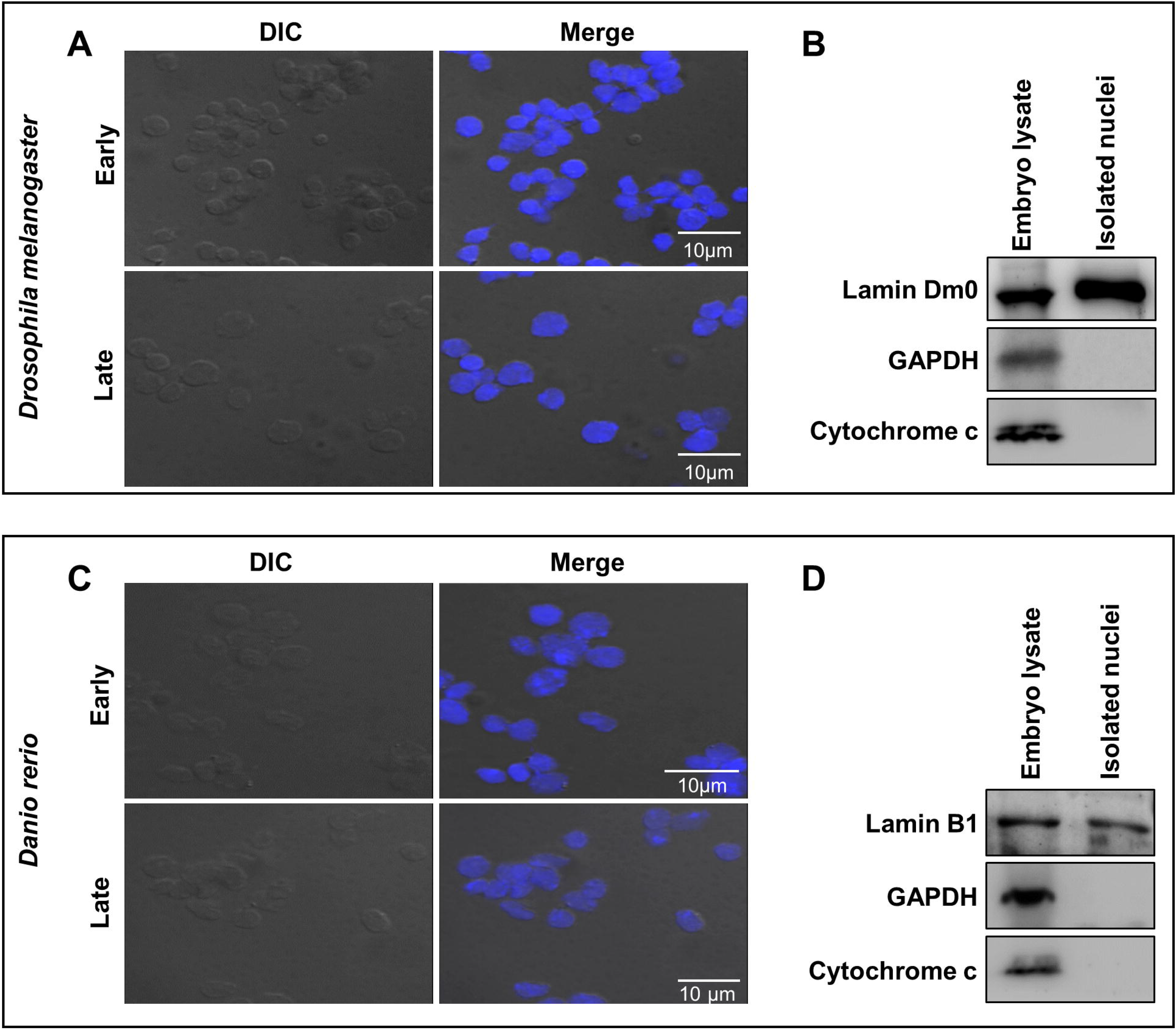
Quality control of nuclei isolated from *D. melanogaste* and *D. rerio* embryos: **(A, C)** DIC images of purified nuclei and those overlapped with DAPI images. The absence of unlyzed cells and membranous contamination indicates the isolation of pure nuclei from both organisms. **(B, D)** Western blots of whole embryo lysate and purified nuclei with antibodies against Lamin, GAPDH and Cytochrome C. Lamin which is a known component of nuclei is present in both whole embryo lysate and nuclei, while GAPDH and Cytochrome C are only present in the former, indicating nuclei preparations are free of cytoplasmic and mitochondrial contamination. Scale Bar: 10 µm.

These nuclei preparations were digested with DNase I and extracted using a sequential extraction method, in 0.4 M NaCl followed by 2 M NaCl, to yield NuMat (Pathak et al. 2007). Aliquots stored at each of the stages were run on an SDS-PAGE gel. The silver stained profiles (Figure 2) of these SDS-PAGE gels showed that NuMat from each developmental stage of both organisms is enriched in high molecular weight proteins. DNase I digestion followed by high salt extraction removes the bulk of proteins including histones from NuMat, which is also evident in western blot analysis with histone H3 antibody. Majority of the histones were extracted in the salt extract fraction thereby leaving the NuMat fractions free of histones and chromatin. On the other hand, most of the Lamin, which is a known component of NuMat, was retained in NuMat.

**Figure 2:**
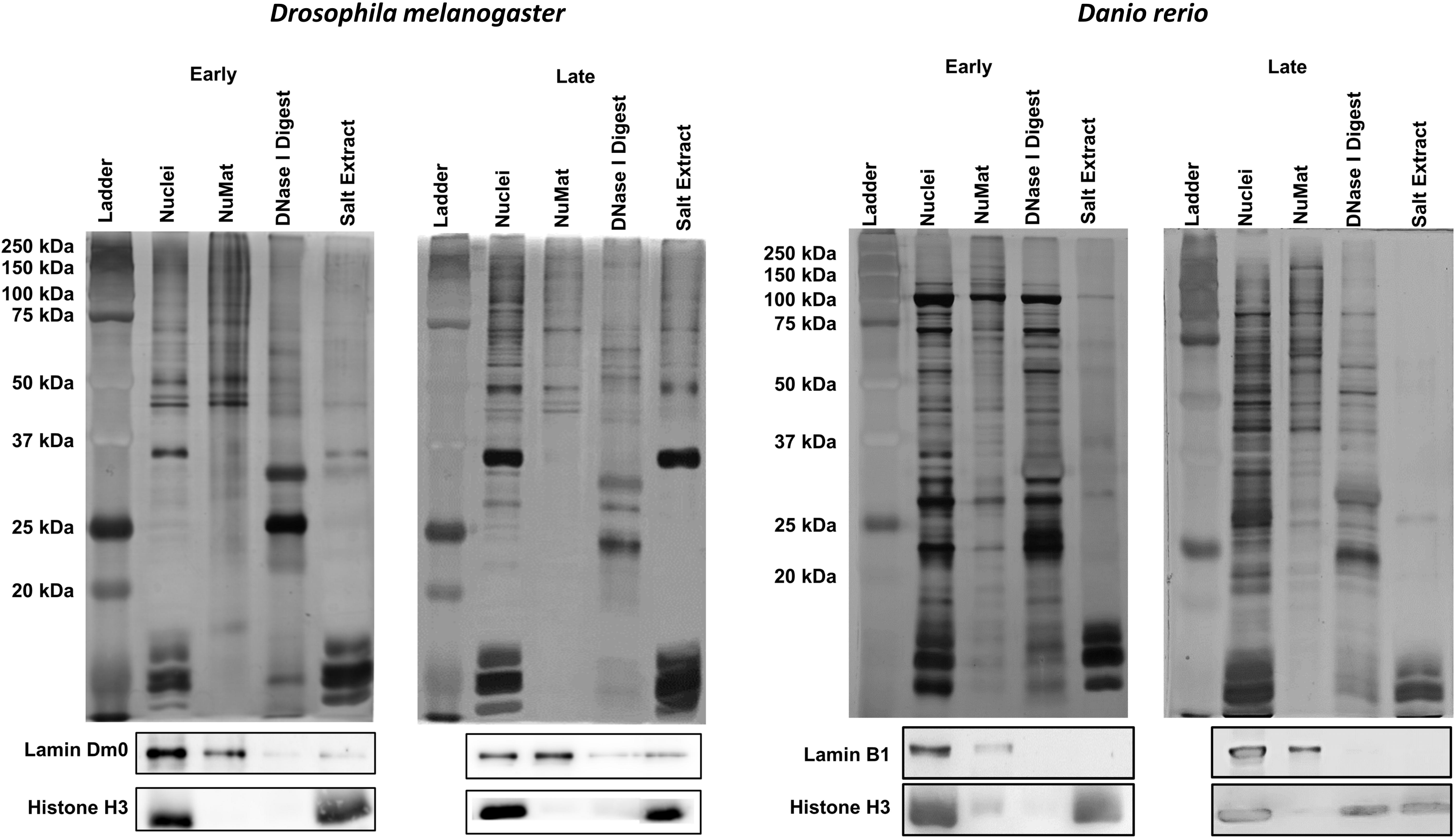
Quality control of NuMat prepared from 0-2 hr (early) and 14-16 hr (late) *D. melanogaster* and 4hpf (early) and 1dpf (late) *D. rerio* embryos: Silver stained profile shows enrichment of high molecular weight proteins in NuMat preparations. Most of the histones are extracted after DNase I digestion and salt extraction in both developmental stages. This is also evident from western blots against histone H3. Western blot against Lamin Dmo0 and Lamin B1 antibody shows that it is present in both nuclei and NuMat of *D. melanogaster* and *D. rerio* respectively.

Estimation of the total molecular composition of nuclei and NuMat showed that while the proportion of DNA, RNA, and proteins in nuclei were 30-38%, 2.3-3%, and 59-67% respectively, it was 6-8%, 7-13%, and 78-86% respectively in NuMat (Figure 3A). Compared to the total molecular composition of the nucleus, NuMat retained ~0.5-0.7% in the form of DNA, 0.7-1.2% in the form of RNA and 7-7.8% in the form of proteins (Figure 3B) in the two organisms.

**Figure 3:**
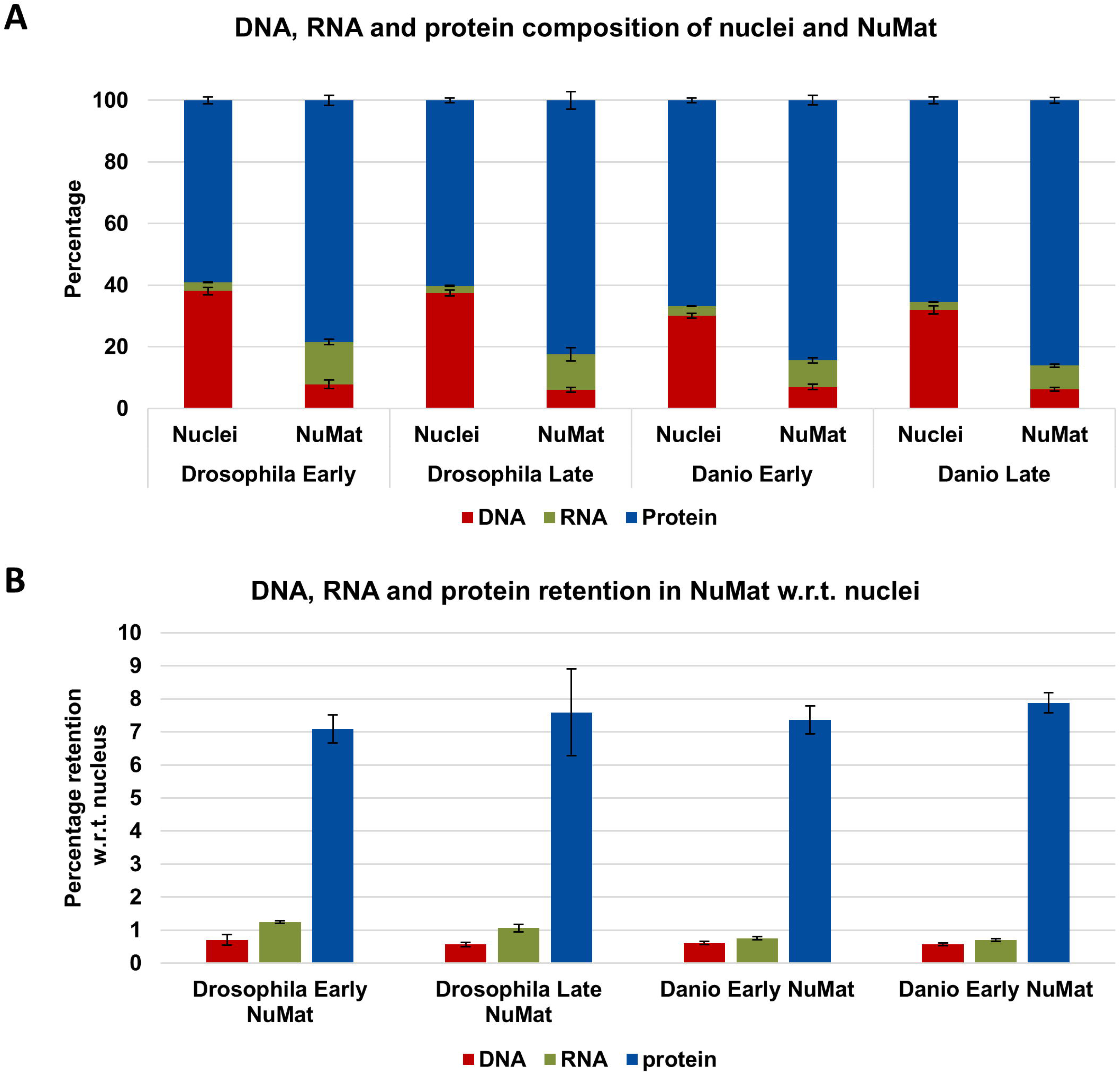
DNA, RNA and protein composition of nuclei and NuMat: **(A)** Percentage composition of DNA, RNA, and protein in nuclei and NuMat. The percentages are calculated using the formula: 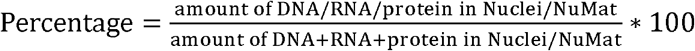. **(B)** Percentage retention of DNA, RNA and proteins in NuMat with respect to the nucleus. The percentages are calculated using the formula: 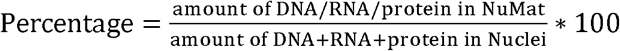. Error bars represent SE.

### Proteome profiling of NuMat by mass spectrometry

In *D. melanogaster*, 1587 and 1572 proteins were identified with high confidence in early and late embryo NuMat respectively (average spearman’s rank correlation of protein intensities between biological replicates was 0.81 and 0.84, respectively, p < 0.0001). Out of these, 1058 were common between the two developmental stages while 529 and 514 were unique to the early and late stage embryo, respectively (Figure 4, Supplementary Table 1). On the other hand, 984 and 1014 high confidence proteins were identified in the early and late embryo NuMat of *D. rerio* (average spearman’s rank correlation of protein intensities between biological replicates was 0.66 and 0.65, respectively, p < 0.0001). 690 of these proteins were common between the early and late embryo NuMat while 294 and 324 were unique to the two stages, respectively (Figure 4, Supplementary Table 2). These 1058 proteins in *D. melanogaster* and 690 proteins in *D. rerio* represent the core NuMat proteins of each species.

**Figure 4:**
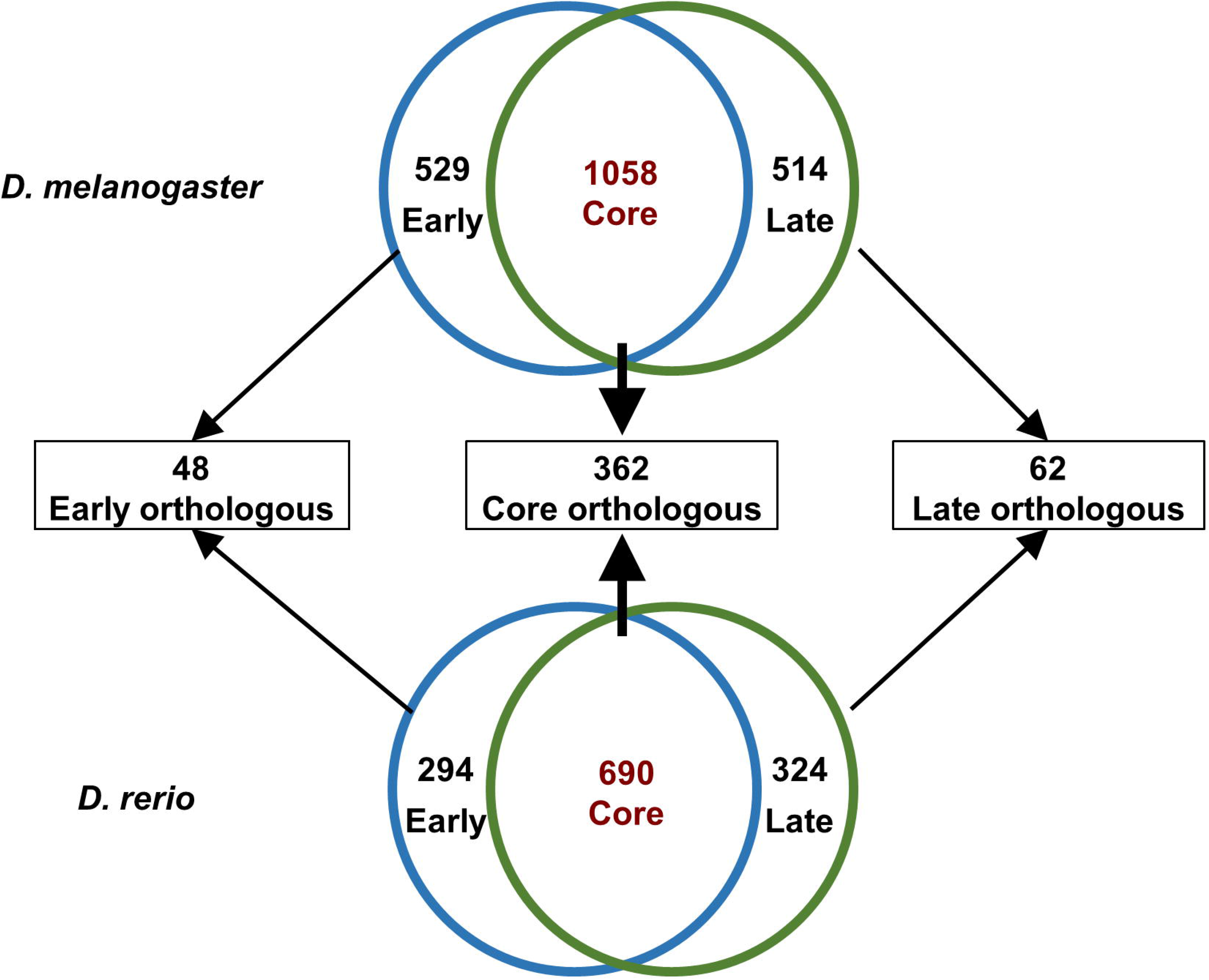
Comparative analysis of *D. melanogaster* and *D. rerio* NuMat proteome: Venn diagram for *D. melanogaster* shows that out of the 1587 and 1572 proteins identified for early and late embryos, 1058 are common between the two developmental stages and represent the core NuMat in *D. melanogaster*. Venn diagram for *D. rerio* shows that out of the 984 and 1014 proteins identified for early and late embryos 690 are common between the two developmental stages and represent the core NuMat in *D. rerio*. Homology search by reciprocal BBH procedure showed that 48 proteins are orthologous among the 529 and 294 unique proteins in the early developmental stages and 62 proteins are orthologous among the 514 and 324 proteins unique to late developmental stages of the two organisms respectively. 362 proteins are orthologous between the core NuMat proteins of the two organisms and are referred to as core orthologous NuMat proteins.

### Comparison of NuMat proteomes of *D. melanogaster* and *D. rerio*

Out of the 529 and 294 proteins unique to NuMat of undifferentiated cells during early development of *D. melanogaster* and *D. rerio* respectively, 48 proteins were orthologous between the two organisms (Figure 4, Supplementary Table 3). These proteins were enriched in cell cycle and DNA replication (Figure 5, yellow bars) related proteins. Similarly, out of the 514 and 324 proteins unique to the NuMat during late development, 62 proteins were orthologous between the two organisms (Figure 4, Supplementary Table 3) and were enriched in transcription factors (Figure 5, green bars). Interestingly, among the 1058 core NuMat proteins of *D. melanogaster* and 690 core NuMat proteins of *D. rerio*, 362 proteins were orthologous (Figure 4, Supplementary Table 3). We refer to these 362 proteins as “core orthologous NuMat proteins”. Functional classification revealed that proteins involved in a wide variety of functions were represented in the core orthologous NuMat proteins but were particularly enriched in RNA binding proteins such as those involved in splicing, RNA metabolism, and transport. Moreover, proteins specific to various nuclear sub-compartments were also exclusively present in core orthologous NuMat proteins (Figure 5, blue bars). We did not find any enriched motifs among the NuMat proteins of either of the organisms.

**Figure 5:**
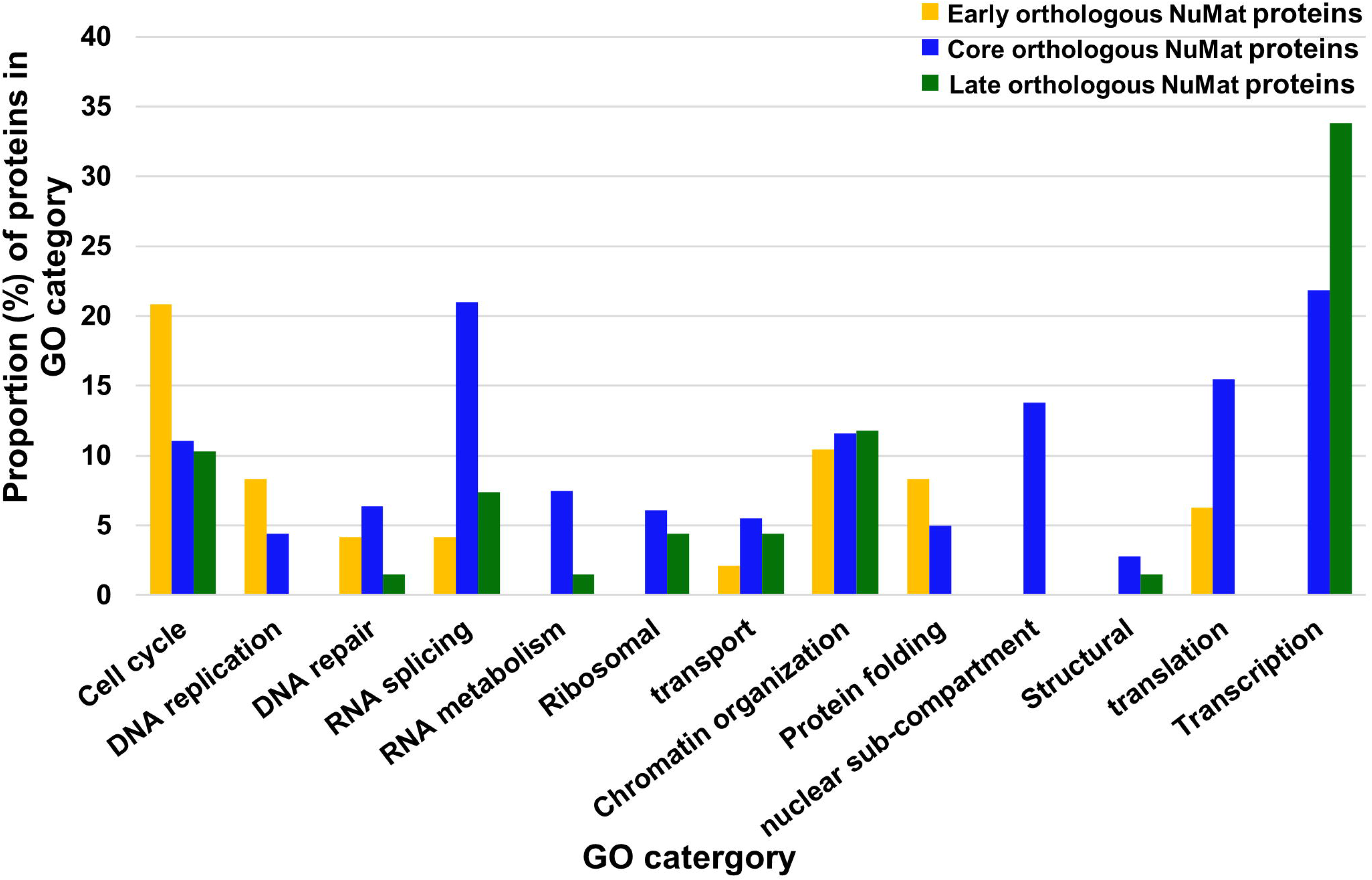
Functional classification of NuMat proteins conserved between *D. melanogaster* and *D. rerio*: Proportion of proteins in early, late and core orthologous NuMat proteins in each functional category (Y-axis) is plotted against different functional categories (X-axis). Orthologous NuMat proteins in the early stages of development are enriched in cell cycle and DNA replication related proteins. The orthologous NuMat proteins in the late stages of development are enriched in transcription factors. Core orthologous NuMat proteins are enriched in RNA splicing, metabolism, transport-related proteins as well as proteins involved in various nuclear subcompartment formation and ribosomal proteins.

### Evolutionary conservation of NuMat proteins across all eukaryotes

In order to identify the core orthologous NuMat proteins which are conserved across eukaryotes, we selected nineteen organisms, based on the completion of their genome assemblies and quality of the gene annotation, with at least one organism from each of the six supergroups (Table 1). Homologs of 362 core orthologous NuMat proteins were searched by the reciprocal BBH procedure in all of the selected eukaryotes. These 362 cores orthologous NuMat proteins, along with the corresponding orthologs in other eukaryotes, were grouped into 358 different protein families. 68 out of 362 core orthologous NuMat proteins had homologs in all selected eukaryotes (Figure 6) and represent the core NuMat proteins which are conserved across eukaryotes. Although proteins from most nuclear functions such as replication, splicing, etc. are represented among these “conserved core NuMat proteins”, they are highly enriched in RNA binding proteins. Interestingly, αTubulin and Actin are also conserved. Out of these 68 proteins 51 proteins had prokaryotic homologs while the other 17 are conserved only among eukaryotes.

**Table 1:** Selected organisms from each eukaryotic supergroup.

| **Eukaryotic Supergroups** | **Organisms** |
| --- | --- |
| Opisthokonts | *Drosophila melanogaster*, *Danio rerio*, *Mus musculus*, *Homo sapiens*, *Saccharomyces cerevisiae* |
| Archaeplastida | *Arabidopsis thaliana*, *Oryza sativa*, *Solanum tuberosum* |
| Amoebozoa | *Dictyostellium discoideum*, *Entamoeba histolytica*,<br>*Acanthamoeba castellanii* |
| Chromalveolates | *Vitrella brassicaformis*, *Toxoplasma gondii*, *Plasmodium falciparum*, *Ectocarpus siliculosus*, *Nannochloropsis gaditana*,<br>*Phaeodactylum tricornutum*, *Thalassiosira pseudonana* |
| Excavates | *Trypanosoma brucei gambiense*, *Trichomonas vaginalis* |
| Rhizaria | *Plasmodiophora brassicae* |

**Figure 6:**
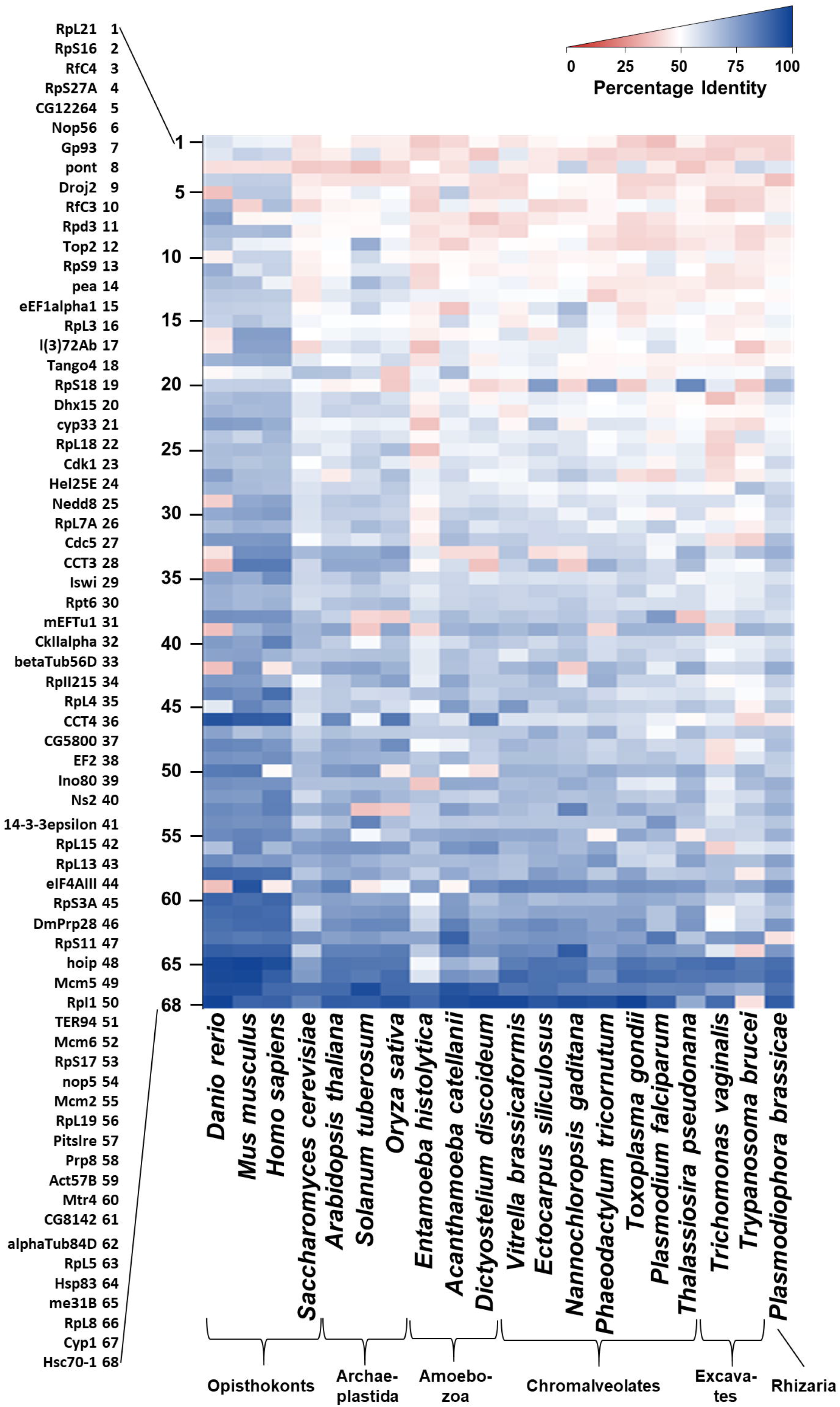
Conserved core NuMat proteins across eukaryotes: Heat map of percentage identity of conserved core NuMat proteins in selected eukaryotes. All identity percentages are with respect to *D. melanogaster* proteins. Rows correspond to gene names for corresponding numbers (Y-axis); columns correspond to selected eukaryotic species(X-axis). Curly brackets (X-axis) indicate the Eukaryotic supergroup to which each organism belongs to. Highlighted gene names represent genes which did not originate from prokaryotes.

### Evolutionary origin of NuMat proteins from prokaryotes

In order to trace the prokaryotic origins of the core NuMat proteins, we used the 362 core orthologous NuMat proteins as seed sequences. We reasoned that since many of the selected eukaryotes occupy very specialized niches and have diverged over 1.5 billion years from each other, many species might have either lost or have diverged versions from the NuMat proteins seen in other extant eukaryotes. 134 out of 362 core orthologous NuMat proteins had homologs with ≥ 30% identity among various prokaryotes. Each of these proteins was added to the corresponding eukaryotic protein family to which these proteins were homologous (132 out of 358 families had homologs). For 226 out of 358 eukaryotic protein families, no homologs could be detected among prokaryotes. Phylogenetic trees were constructed for each protein family that had prokaryotic homologs. The prokaryotic proteins were grouped according to their respective phyla, and their relationship with the eukaryotic proteins was plotted (Figure 7). For 43 protein groups, a sister group relationship could be established between eukaryotes and various archaeal phylum, out of which 27 belonged to Euryarchaeaota. Interestingly, except RpL8 (60S Ribosomal protein L8) all other eukaryotic ribosomal proteins have a sister group relationship with archaea. The other overrepresented phylum having a sister group relationship with eukaryotic proteins belonged to proteobacteria with 21 proteins. Therefore, out of the 134 core orthologous NuMat proteins which have homologs in prokaryotes, a large fraction have a sister group relationship with proteins in Euryarchaeota and Proteobacteria, the two most prominent superphyla in this analysis. For 48 protein groups, no sister group relationship could be established with eukaryotic proteins.

**Figure 7:**
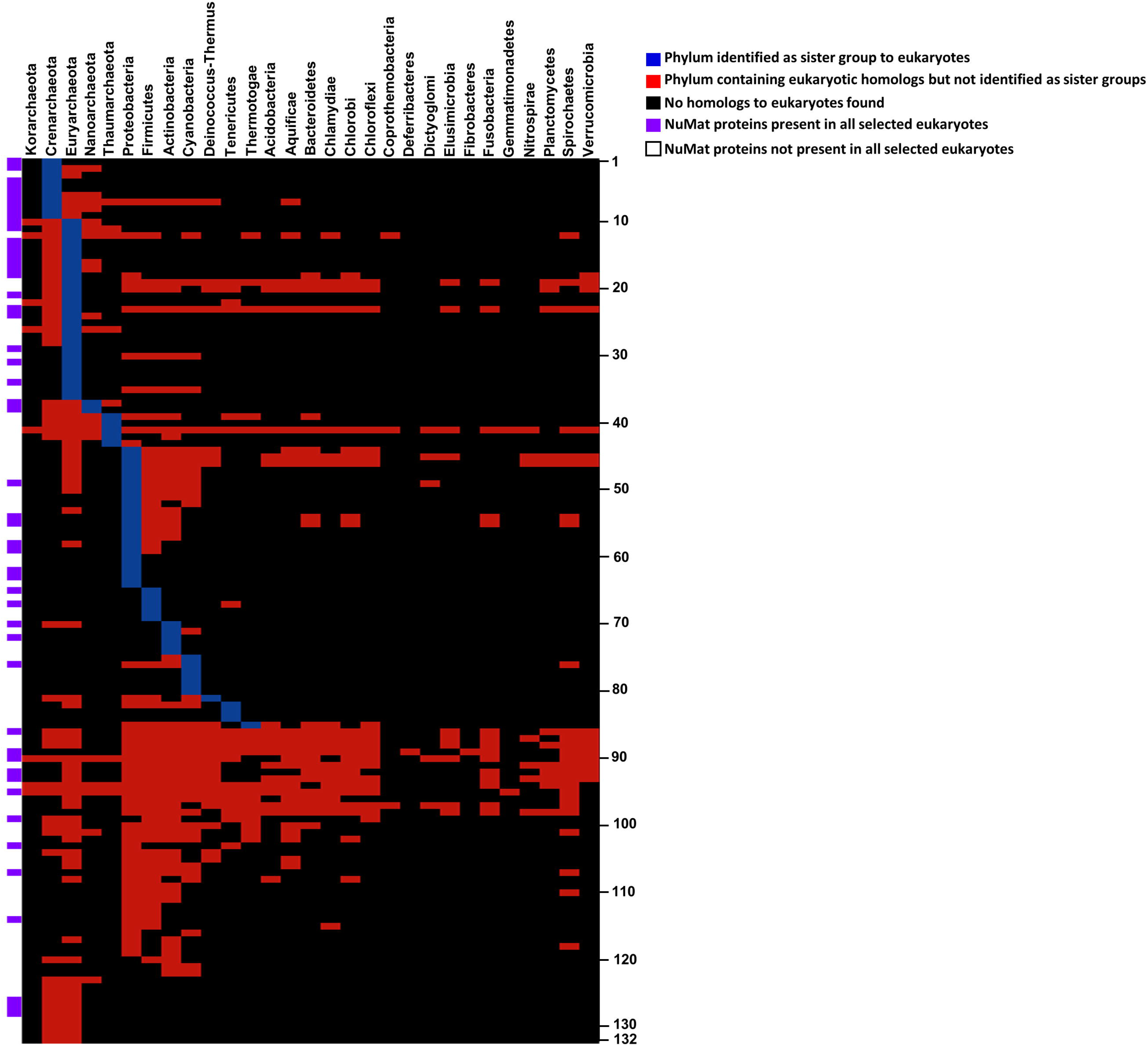
Bacterial taxonomic groups found to have homologs of core orthologous NuMat proteins: Rows correspond to 132 eukaryotic gene families (gene names correspond to *D. melanogaster* orthologs) found to have orthologs in prokaryotes; columns correspond to 29 prokaryotic phyla. Gene names corresponding to the respective numbers of each row are shown in Table 2. Blue cells represent the gene (row) for which a sister group relationship could be established between the selected eukaryotes and organisms from the corresponding prokaryotic phyla (column). Red cells represent genes for which no sister group relationship could be established. Purple cells represent genes which are also conserved across all selected eukaryotes. The same is indicated by ‘*’ in Table 2.

## Discussion

Ever since its discovery, the NuMat allowed the conceptualization of a structural platform for various nuclear functions and which could be considered responsible for the structural integrity of the nucleus (Stein et al. 2003; Zink et al. 2004). Since proteins are the major structural constituents of NuMat, many studies have strived to identify its core proteome. Most of these studies, with few exceptions (Kallappagoudar et al. 2010; Varma and Mishra 2011), are based on cell lines (Albrethsen et al. 2010; Engelke et al. 2014) which does not reflect the *in vivo* state accurately (Antoni et al. 2015). In the present study, we used the developing embryos of *D. melanogaster* and *D. rerio* as model organisms. We found that ~50% of the NuMat proteins identified in the two organisms are preserved in both differentiated and undifferentiated cells (Figure 4). These proteins represent the core NuMat proteome, as they are associated with the NuMat, irrespective of the developmental stage. On the other hand, proteins unique to a developmental stage represent the cell type-specific additions. It is possible that some of the identified NuMat proteins are contaminants. However, such contaminants are likely to be stochastic and specific to the organisms. By comparing the NuMat proteomes of two evolutionarily distant organisms, we eliminated such proteins from the orthologous NuMat proteins. We found 362 proteins that are conserved between the core NuMat proteins of fish and fly, which implies that these proteins are not only present in NuMat irrespective of the cell type but also irrespective of the organism. Moreover, homologs of 262 of these proteins were also found in the NuMat (NaCl extracted fraction) of PD36 pre-B cell lines in a previous study (Engelke et al. 2014) suggesting their authenticity and importance in NuMat. Our finding that 48 and 62 proteins are conserved in the early and late embryo NuMat preparations, respectively of the two evolutionarily distant organisms indicates their importance in nuclear structure during embryonic development.

Many of the core orthologous NuMat proteins are involved in diverse nuclear functions, some of which occur at different stages of the cell cycle. This suggests that all of the 362 core orthologous proteins do not associate with the NuMat at the same time, but, instead, associate during different stages of the cell cycle, implying that the NuMat is not a static structure, but is dynamic as per the changing requirements of the cell (Roix and Misteli 2002). This is also reflected in the early and late stage conserved proteins. For instance, the homologous proteins of the early NuMat are enriched in cell cycle, and DNA replication proteins (Figure 5) since blastula cells are known to divide very fast (Gilbert 2000). On the other hand, the homologous proteins in the late NuMat are enriched in transcription factors which are required for the functions of the vast variety of cells populating the late embryo. The dynamic nature of NuMat was also exemplified by a recent study which showed that during mitosis, 67% of the NuMat proteome is packaged along with chromosomes in the form of mitotic chromosome scaffold during mitosis and helps in transfer of architectural information from mother to daughter nuclei (Sureka et al. 2018).

Interestingly, 132 protein families out of 358 core orthologous NuMat protein families had prokaryotic homologs, indicating that pre-existing prokaryotic versions of these proteins were exapted for nuclear architecture in eukaryotes. Notably, many ribosomal proteins, present in the core orthologous NuMat, were found to have originated from archaea (Figure 7, Table 2). Since transcription and translation are coupled in prokaryotes, it is possible that nuclear translation by ribosomal proteins was retained during the early stages of evolution of the nucleus, and was later adopted as a proofreading mechanism for mRNAs, as was suggested by studies on nonsense mediated decay in the nucleus (Iborra et al. 2004; David et al. 2012; Reid and Nicchitta 2012). This idea is further supported by a recent proteomic analysis which revealed the putative nuclear localization of all ribosomal proteins in *Arabidopsis thaliana* (Palm et al. 2016). Similarly, the NuMat associated replication proteins seem to have originated from archaea. On the other hand, since it is well accepted that mitochondria is involved in various stress response pathways and that it evolved from α-proteobacterial endosymbiont, the observation that many stress response proteins (such as DmRH19, DmRH24, bel, me31B) and those involved in protein folding (such as Gp93, Hsp83, and Hsc70), found in core NuMat, are most closely related to proteobacteria, suggests that the NuMat proteins of proteobacterial origin were exapted for nuclear functions following the endosymbiosis of the mitochondrial ancestor (Figure 7, Table 2) (Margulis 1970; Knowlton and Salfity 1996; Gray 1999; Suhm and Ott 2017). The fact that seven of the proteins originating from proteobacteria were also conserved across all the eukaryotic supergroups, also provides evidence to support the hypothesis that the nucleus evolved after the endosymbiosis of proteobacteria (Martin and Koonin 2006; Dacks et al. 2016). We also noticed that many different prokaryotes contributed different protein families which constitute the NuMat, thus imploring us to consider the role of horizontal gene transfer (HGT) in spreading genes among extant prokaryotic communities.

**Table 2:** List of 132 eukaryotic gene families of core orthologous NuMat proteins having prokaryotic homologs.

| No. | Gene<sup>Ⓜ</sup> | No. | Gene | No. | Gene | No. | Gene |
| --- | --- | --- | --- | --- | --- | --- | --- |
| 1 | Mcm2* | 35 | eIF4A | 69 | pit | 103 | Top2* |
| 2 | RpL13A* | 36 | RpL24 | 70 | Mtr4* | 104 | Aldh |
| 3 | CCT8 | 37 | Hoip* | 71 | Rack1 | 105 | Scsalpha |
| 4 | RpS27* | 38 | RpL4* | 72 | I(3)72Ab* | 106 | PCB |
| 5 | Mcm5* | 39 | RpS2 | 73 | alphaCOP | 107 | Cyp1* |
| 6 | RfC3* | 40 | RpS13 | 74 | obe | 108 | SMC1 |
| 7 | eEF1alpha1* | 41 | RpL23 | 75 | Taf5 | 109 | CG12262 |
| 8 | CG8142* | 42 | RpS23 | 76 | cyp33* | 110 | Nc73EF |
| 9 | Mcm6* | 43 | CG8939 | 77 | Q9VT19 | 111 | Arc42 |
| 10 | RpL5* | 44 | ND-30 | 78 | Caper | 112 | G5599 |
| 11 | RpL7A* | 45 | Hsc70-5 | 79 | U4-U6-60K | 113 | Cdk2 |
| 12 | RpS14a | 46 | Hsc70-3 | 80 | CstF64 | 114 | CG5800* |
| 13 | RpS17* | 47 | mahe | 81 | ldh3b | 115 | prtp |
| 14 | RpL19* | 48 | Dbp73D | 82 | CG9143 | 116 | SMC3 |
| 15 | RpS18* | 49 | CG12264* | 83 | kis | 117 | mask |
| 16 | RfC4* | 50 | DmRH19 | 84 | dom | 118 | ERp60 |
| 17 | RpS11* | 51 | Dbp45A | 85 | Gnf1 | 119 | Idh |
| 18 | RpS16* | 52 | bel | 86 | TER94* | 120 | CCT2 |
| 19 | Rpt2 | 53 | Ddx1 | 87 | Rpl1* | 121 | dSki8 |
| 20 | RpII140 | 54 | Gp93* | 88 | smid | 122 | wds |
| 21 | CCT3* | 55 | Hsp83* | 89 | mEFTu1* | 123 | RpS8 |
| 22 | sta | 56 | alpha-KGDHC | 90 | RpL8* | 124 | Ski6 |
| 23 | Rpt6* | 57 | Hlc | 91 | Hsc70-4 | 125 | RpII33 |
| 24 | RpL15* | 58 | Hel25E* | 92 | Droj2* | 126 | RpL3* |
| 25 | RpS19a | 59 | me31B* | 93 | Hsc70-1* | 127 | RpL21* |
| 26 | RpS9* | 60 | DHX8 | 94 | waw | 128 | CCT4* |
| 27 | RfC38 | 61 | CG6994 | 95 | EF2* | 129 | RpS6 |
| 28 | CG3756 | 62 | Act57B* | 96 | Gapdh2 | 130 | RpL34a |
| 29 | Rpd3* | 63 | Dhx15* | 97 | Lon | 131 | RpL24-like |
| 30 | CG7483 | 64 | cg5589 | 98 | blw | 132 | RpS4 |
| 31 | nop5* | 65 | lswi* | 99 | RpII215* |  |  |
| 32 | RpS3 | 66 | Mi-2 | 100 | I(1)G0156 |  |  |
| 33 | Non1 | 67 | Ino80* | 101 | Jafrac1 |  |  |
| 34 | Nop56* | 68 | Chd1 | 102 | CG5028 |  |  |

Genes marked with ‘\*’ are also conserved across eukaryotes.
<sup>Ⓜ</sup> Gene names correspond to *D. melanogaster* orthologs

Among the 132 protein families of the core orthologous NuMat that have a prokaryotic origin, only 51 (Figure 7, Table 2) were conserved across all selected eukaryotes. Since eukaryotes do not acquire genes through continual HGT like prokaryotes do (Ku and Martin 2016), this may indicate the loss or evolutionary divergence of certain proteins under a variety of selection pressures in some organisms but still relevant for nuclear functions in other lineages. Nonetheless, 68 out of the 362 core orthologous NuMat proteins were conserved across all eukaryotic supergroups, pointing towards their ancestral nature and continued role in NuMat (Figure 6). However, the 17 proteins which did not have prokaryotic homologs, were later additions specific to the changing requirements of emerging nuclear functions, such as those involved in splicing (Interestingly 6 out of these 17 proteins are invovled in splicing) (Martin and Koonin 2006). However, since they are conserved among all eukaryotes they are most likely to have evolved before the emergence of the LECA. In contrast to the 83 protein families which diverged in different eukaryotic lineages, these 68 proteins have not diverged much despite 1.5 billion years of evolution, endorsing their universal importance in nuclear structure and function. Intriguingly, 26 and 57 out of these conserved core NuMat proteins were also found to be essential in yeast and humans, respectively, further supporting the importance of these proteins (Giaever et al. 2002; Blomen et al. 2015). Additionally, these 68 proteins are more identical among all multicellular eukaryotes used in our study, than with unicellular eukaryotes, pointing towards a correlation between complexity of the organism and NuMat proteome. It is also possible that many of the 68 conserved core NuMat proteins do not localize to the NuMat in some of the eukaryotes. Our approach, which involves identification of the NuMat proteins from two multicellular organisms belonging to the opisthokonts supergroup, and then using a bioinformatic approach to identify potential NuMat proteins in other supergroups may lead to false positive identifications and cannot be a substitute for experimental identification of NuMat proteins from each of the supergroups, especially the unicellular eukaryotes. However, since all of these 68 proteins were experimentally found to be present in both *D. melanogaster* and *D. rerio*, they most likely are a part of the core NuMat at least in metazoans.

In conclusion, we identified the core NuMat proteins in *D. melanogaster* and *D. rerio* by comparing the respective proteomes from the different types of cells present in developing embryos of the two distantly related organisms. This minimum set of proteins, the core NuMat proteome, allowed us to explore the early stage of nuclear evolution, and, by extension, that of eukaryotes. We find that 132 protein families form the initial substrate for nuclear architecture by recruiting existing prokaryotic proteins for this purpose. While a subset of these, 51 proteins, were retained as the conserved core NuMat proteins, others were either lost or diverged to group specific specializations. In this process additional (~200) proteins were recruited for various aspects of nuclear structure and function in metazoans. This study, therefore, provides an important resource for future studies on nuclear architecture and function.

## Materials and Methods

### Embryo Collection

#### *Drosophila melanogaster* (Fruit fly)

For all the experiments, Canton-S (CS) flies were used. Plates containing solidified fly food (2.5% w/v sucrose, 25% v/v grape juice, 2.5 % w/v agar, 0.5% v/v benzoate) streaked with yeast paste were placed inside large cages populated with adult flies. The fly cages were first synchronized by changing fly food plates every hour, for three hours. For collecting 0-2 hr embryos, the fly food plates were incubated in such synchronized cages for two hours before removing and collecting the embryos. For collecting 14-16 hr old embryos, plates were removed after two hours from synchronized cages and incubated at 25° C for an additional 14 hrs before collecting the embryos.

Immediately after collection, the embryos were dechorionated with 50% Chlorex (4-6% sodium hypochlorite) for 5 minutes with continuous stirring. The removal of chorion was assessed by the floating chorion on top of the solution. The dechorionated embryos were washed thoroughly with running water using a mesh cloth to completely remove Chlorex. These embryos were then dried and weighed before use for NuMat preparation.

#### *Danio rerio* (Zebrafish)

In-house bred adult *D. rerio* were maintained according to the standard procedures. Males and females were separated by a wire mesh and kept at 28° C overnight, in the dark. In the morning, the separating wire mesh was removed and the males and females were put in a tank over a mesh large enough to allow embryos to pass through. The lights were switched on and fishes allowed to lay eggs for 1 hr. The embryos were then collected and washed with running water over fine mesh before transferring to embryo medium (13.6 mM NaCl, 0.5 mM KCl, 0.0025 mM Na_2_HPO_4_, 0.0044 mM KH_2_PO_4_, 0.0064 mM CaCl_2_, 0.005 mM MgSO_4_, 0.0041 mM NaHCO_3_) and incubated at 28° C. For 4 hpf (hours post fertilization) embryos, the embryos were removed from incubator after 3 hrs and dechorionated. For 1dpf (days post fertilization) embryos, the embryos were removed from the incubator after 23 hrs and dechorionated.

Embryos were dechorionated by incubating in 2 mg/ml Pronase in embryo medium for 5 mins at 28° C and then washed with embryo medium with gentle agitation so that the chorion floats away from the embryos. The washing was repeated several times. The dechorionated embryos were then used for NuMat preparation.

### Nuclei Isolation

#### *D. melanogaster* embryos

Dechorionated *D. melanogaster* embryos were chilled on ice and homogenized in pre-chilled DmNIBI (*D. melanogaster* nuclear isolation buffer I) (15 mM Tris pH 7.4, 0.25 M sucrose, 40 mM KCl, 1 mM EDTA pH 8.0, 0.1 mM EGTA, 0.25 mM Spermidine, 0.1 mM Spermine, 0.1 mM PMSF and 0.1 μM Aprotinin) using a Dounce homogenizer (10 strokes, 400 rpm) and filtered through two layers of Mira cloth (Calbiochem). The filtrate was centrifuged at 600g at 4° C for 2 min to remove crud and unbroken cells. The resulting supernatant was mixed gently with an equal volume of DmNIBII (*D. melanogaster* nuclear isolation buffer II) (15 mM Tris pH 7.4, 1.8 M sucrose, 40 mM KCl, 1 mM EDTA pH 8.0, 0.1 mM EGTA, 0.25 mM Spermidine, 0.1 mM Spermine, 0.1 mM PMSF and 0.1 μM Aprotinin) and centrifuged at 6000g at 4° C to isolate the nuclear pellet. The pellet obtained was allowed to swell in DmNIBI for 5 min at 4° C, and then washed twice by resuspending it completely in DmNIBI and centrifuging at 1000g for 10 min at 4° C. The nuclear pellet thus obtained was suspended in 2 ml of DmNIBI.

#### *D. rerio* embryos

Dechorionated *D. rerio* embryos were chilled on ice and homogenized in pre-chilled DrNIBI (*D. rerio* nuclear isolation buffer I) (10 mM HEPES pH 7.4, 0.25 M sucrose, 40 mM KCl, 5 mM MgCl_2_, 0.25 mM Spermidine, 0.1 mM Spermine, 0.1 mM PMSF and 0.1 μM Aprotinin) using a Dounce homogenizer (5 strokes, 300 rpm) and filtered through two layers of Mira cloth (Catalog# 475855, Calbiochem). The filtrate was centrifuged at 200g at 4° C for 2 min to remove crud and unbroken cells. The resulting supernatant was layered over of DrNIBII (*D. rerio* nuclear isolation buffer II) (10 mM HEPES pH 7.4, 1.25 M sucrose, 40 mM KCl, 5 mM MgCl_2_, 0.25 mM Spermidine, 0.1 mM Spermine, 0.1 mM PMSF and 0.1 μM Aprotinin) and centrifuged at 6000g at 4° C to isolate the nuclear pellet. The pellet obtained was allowed to swell in DrNIBI for 5 min at 4° C, and then washed twice by resuspending it completely in DrNIBI and spinning at 1000g for 10 min at 4° C. The nuclear pellet thus obtained was suspended in 2 ml of DrNIBI.

The amount of nuclei obtained was quantified by lysing 2 µl nuclear suspension in 0.5 % SDS and taking OD reading at 260 nm. The nuclear pellet was then resuspended at a concentration of 10 ODU/ml in (Dm/Dr)NIBI and stabilized by incubating at 37° C for 20 mins before centrifuging to recover the nuclei.

### Preparation of NuMat

10 ODUs of nuclei from *D. melanogaster* or *D. rerio* were incubated in digestion buffer (DB) (DB - 20 mM Tris, pH 7.4, 20 mM KCl, 70 mM NaCl, 10 mM MgCl_2_, 0.125 mM spermidine, 0.05 mM spermine, 0.1 mM PMSF, 0.5% Triton X-100) with 100 µg/ml DNase I at 4° C for 1 hr. Digestion was followed by extraction in extraction buffer (10 mM HEPES, pH 7.5, 4 mM EDTA, 0.25 mM spermidine, 0.1 mM PMSF, 0.5% Triton X-100) containing 0.4 M NaCl, with gentle rocking, for 5 min at 4° C, followed by extraction in extraction buffer containing 2 M NaCl for an additional 5 minutes to obtain NuMat (Pathak et al. 2007). The NuMat pellet was washed with wash buffer (5 mM Tris, 20 mM KCl, 1 mM EDTA, 0.25 mM spermidine, 0.1 mM PMSF) twice and used for further processing.

### SDS-PAGE, western blotting

Samples were resolved on 12% SDS-PAGE gel and were either silver stained or transferred to PVDF membrane (Millipore IPVH00010). The membranes were probed with Lamin Dm0 (DSHB ADL101) (for *D. melanogaster*) or Lamin B1(abcam ab16048) (*for D. rerio*), histone H3 (abcam ab71956), GAPDH (abcam ab8245) and cytochrome C (abcam Ab13575) antibody.

### In-gel digestion and protein identification by LC-MS/MS

For tryptic digestion, 60 µg of protein were resolved on 12% SDS-PAGE. The gel was cut into four pieces, sliced and dehydrated twice using 50% ACN followed by 100% ACN and the proteins digested with ~100 µl of Trypsin solution (10 µg/ml, Promega) in 25 mM ammonium bicarbonate by incubating overnight at 37° C. Digested peptides were eluted with 50% ACN and 5% TFA, desalted and loaded on to the mass spectrometer.

Tryptic peptides were loaded on C18 columns (5 µm, 75 µm×100 mm) (BIOBASIC product#72105-107582). The peptides were separated using gradients ranging from 0 - 95 % acetonitrile, followed by washes with 95 % acetonitrile. The length of the gradient was adjusted to 140 mins. The Thermo Easy nLC II system was directly connected with Thermo fisher scientific Q-exactive instrument using a nano-electrospray source. The nano-source was operated at 1.8 – 2.2 kV and the ion transfer tube at 285-300° C without sheath gas. The mass spectrometer was operated in a data-dependent mode. The scans were acquired with resolution 70,000 at m/z 200-2000 in Orbitrap mass analyser with lock mass option enabled for the 445.120024 ion. The 10 most intense peaks containing double or higher charge state were selected for sequencing and fragmented in the orbitrap by using HCD with a normalized collision energy of 28-30%. Dynamic exclusion was activated for all sequencing events to reduce the chances of repeated sequencing. Peaks selected for fragmentation more than once within 30 s were excluded from selection.

### Mass spectrometric data analysis

The raw spectra obtained were processed using MaxQuant (version 1.5.5.1) with associated Andromeda search engine. Search was performed against *D. melanogaster* and *D. rerio* proteome downloaded from UniProtKB. Trypsin/P was specified as the cleavage enzyme. The search included oxidation of methionine and N-terminal acetylation as variable modifications. Default values of 1% false discovery rate (FDR) were used for PSM and protein identification. For all other parameters default options in MaxQuant were used. Proteins identified with multiple peptides and at least one unique peptide at <1%FDR were used for analysis. The mass spectrometry proteomics data have been deposited to the ProteomeXchange Consortium via the PRIDE partner repository with the dataset identifier PXD012926 (Perez-Riverol et al. 2019). It can be accessed using the username ‘’ and password ‘uQwPXBUA’.

Three biological replicates from independent biological preparations of NuMat were analyzed for each developmental stage from both organisms. Only the proteins which were identified in all three biological replicates with at least two unique peptides were accepted as true NuMat components and were used for further analysis. Reliability of quantification measurements between biological replicates was tested using Spearman Rank correlation. Functional classification of proteins was done manually based on Gene Ontology annotations on UniprotKB.

### Bioinformatic analysis of evolutionary conservation of NuMat proteins

The reciprocal best BLAST hit (BBH) procedure (Tatusov 1997) was used to identify orthologs between the proteins of *D. melanogaster* and *D. rerio*. Proteins were considered orthologous if they had ≥35% identity and an e-value of ≤ 1 × 10^−10^. Proteomes of nineteen eukaryotic organisms, with at least one organism from each of the six supergroups of eukaryotes (Table 1), was downloaded from UniProtKB. Homologs of conserved NuMat proteins in these organisms were identified using BBH procedure with the aforementioned identity and e-value cut-offs. Proteins were grouped into families using MCL with an expansion value of 30 (Enright 2002).

In order to identify prokaryotic homologs of NuMat proteins, the reciprocal BBHs procedure was used against the Swiss-Prot prokaryotic proteome database, and proteins were considered orthologous only if they had ≥30% identity and an e-value of ≤ 1 × 10^−10^. Orthologous prokaryotic proteins were added to protein families (generated by MCL) using an in-house script. The proteins of each family were aligned using MAFFT (version 7.407) (Katoh and Standley 2013) and the phylogenetic trees calculated using RAxML (version 8) using the WAG substitution model (Stamatakis 2014). No additional evaluation or optimization of trees was done beyond what is performed by default in RAxML. The best trees were rendered using ete3 (Huerta-Cepas et al. 2016). The trees were manually analysed for sister group relationships between all eukaryotes and prokaryotic phyla.

## Acknowledgements

We would like to thank Dr. Rashmi U. Pathak and Dr. Divya Tej Sowpati for valuable discussions and suggestions, and Dr. Swasti Raychaudhuri, Dr. Suman S. Thakur and Y. Kameshwari for their help in proteomics analysis.

